# Nanobodies against SARS-CoV-2 non-structural protein Nsp9 inhibit viral replication by targeting innate immunity

**DOI:** 10.1101/2023.10.12.561992

**Authors:** Tomas Venit, Jeremy Blavier, Sibusiso B. Maseko, Sam Shu, Lilia Espada, Christopher Breunig, Hans-Peter Holthoff, Sabrina C. Desbordes, Martin Lohse, Gennaro Esposito, Jean-Claude Twizere, Piergiorgio Percipalle

## Abstract

Nanobodies are emerging as critical tools for drug design. Several have been recently created to serve as inhibitors of SARS-Cov-2 entry in the host cell by targeting surface-exposed Spike protein. However, due to the high frequency of mutations that affect Spike, these nanobodies may not target it to their full potential and as a consequence, inhibition of viral entry may not be efficient. Here we have established a pipeline that instead targets highly conserved viral proteins that are made only after viral entry into the host cell when the SARS-Cov-2 RNA-based genome is translated. As proof of principle, we designed nanobodies against the SARS-CoV-2 non-structural protein Nsp9, required for viral genome replication. To find out if this strategy efficiently blocks viral replication, one of these anti-Nsp9 nanobodies, 2NSP23, previously characterized using immunoassays and NMR spectroscopy for epitope mapping, was encapsulated into lipid nanoparticles (LNP) as mRNA. We show that this nanobody, hereby referred to as LNP-mRNA- 2NSP23, is internalized and translated in HEK293 cells. We next infected HEK293-ACE2 cells with multiple SARS-CoV-2 variants and subjected them to LNP-mRNA-2NSP23 treatment. Analysis of total RNA isolated from infected cells treated or untreated with LNP-mRNA-2NSP23 using qPCR and RNA deep sequencing shows that the LNP-mRNA-2NSP23 nanobody protects HEK293-ACE2 cells and suppresses replication of several SARS-CoV-2 variants. These observations indicate that following translation, the nanobody 2NSP23 inhibits viral replication by targeting Nsp9 in living cells. We speculate that LNP-mRNA-2NSP23 may be translated into an innovative technology to generate novel antiviral drugs highly efficient across coronaviruses.

## Introduction

Coronaviruses that infect humans (HCoV) can be classified based on their pathogenicity into seasonal and highly pathogenic viruses. Seasonal viruses including HCoV-229E (Hamre and Procknow, 1966), HCoV-OC43 (McIntosh et al., 1970), HCoV-NL63 (van der Hoek et al., 2004), and HCoV-HKU1 (Woo et al., 2005), induce mild upper-respiratory-tract symptoms responsible for up to 30% of common colds in adults but in infants, the elderly, and immunocompromised individuals, are able to cause a severe lower-respiratory tract disease (Matoba et al., 2015; Otieno et al., 2022). The second category comprises severe acute respiratory syndrome virus (SARS)-CoV and -CoV-2, as well as Middle East respiratory syndrome (MERS)-CoV, which can rapidly spread from the upper-respiratory epithelial cells, infect the lower-respiratory tract and induce severe diseases (Zhong et al., 2003). These viruses can cause epidemics and global pandemics such as COVID-19 (V’Kovski et al., 2021; Wang et al., 2020). Coronaviruses, similarly to other viruses, have the potential to mutate and recombine to generate new variants that could escape current available vaccines. It is thus of utmost importance to develop novel antiviral molecules targeting conserved processes of coronavirus replication, such as the conserved replication-transcription complex (RTC). The precise strategy used by coronaviruses to effectively replicate their large genome is not fully understood (Brian and Baric, 2005; Woo et al., 2010), but the stability and proper organization of all components of coronaviruses-encoded RTC, including the RNA-dependent RNA polymerase (RdRp) and associated factors, play crucial roles in viral mRNAs synthesis from their RNA templates (Brian and Baric, 2005; Fehr and Perlman, 2015). Structural snapshots of the SARS-CoV-2 RTC have been reported at atomic resolution. The mini RTC complex is assembled by a number of non-structural proteins (Nsp) including Nsp7-2xNsp8-Nsp12-2xNsp13- and the RNA-binding protein, Nsp9 necessary for RTC function (Yan et al., 2020). Although Nsp9 has a strong tendency to oligomerize (Ponnusamy et al., 2008; Zhang et al., 2020), within the RTC, it is primarily in a monomer or homodimer state (Yan et al., 2021). Previous evidence linking Nsp9 dimerization and viral propagation (Miknis et al., 2009; Sutton et al., 2004; Zeng et al., 2018) is well explained by the coincidence of all Nsp9 dimerization interface residues with its binding contacts in the catalytic centre of the RNA-dependent RNA polymerase Nsp12 (Yan et al., 2021). Even the higher affinity for single stranded RNA of Nsp9 dimers compared to monomers (Egloff et al., 2004; Zeng et al., 2018) did not support a functional homodimeric state because of the relative weakness of the binding (Littler et al., 2020) suggesting the occurrence of a replication complex (Miknis et al., 2009) well ahead the actual observation. Due to its unique role in viral RNA transcription, Nsp9 protein could serve as a compelling target for inhibiting coronaviruses replication.

Nanobodies have emerged as ideal tools for a drug design (Hamers-Casterman et al., 1993) and several have been made available against the surface-exposed Spike protein to primarily block viral entry in the host cell (Dong et al., 2020; Huo et al., 2020; Ma et al., 2022; Maeda et al., 2022; Wrapp et al., 2020). However, given the frequency of mutations that affect Spike, the efficiency of these nanobodies as potential antivirals is questionable. We, therefore, reasoned that the conserved Nsp9 would be a more efficient target to design novel compounds that robustly inhibit Sars-CoV-2 replication with the potential to work as pan-coronavirus antivirals. We generated 136 unique nanobodies against Nsp9 and the most promising ones, namely 2NSP23 and 2NSP90, were expressed, purified and characterized (Esposito et al., 2021). The results from immunoassays show that these nanobodies effectively and specifically recognize viral Nsp9 in saliva samples from COVID-19 patients. NMR analysis and molecular dynamics simulations identified the epitopes on the wild-type Nsp9 protein recognized by the anti-NSP9 nanobodies and revealed that nanobody treatment lead to predominant non-functional tetrameric assembly of Nsp9 (Esposito et al., 2021; Esposito and Percipalle, 2023, personal communication).

In the present study, we therefore investigated if nanobody 2NSP23 inhibits Sars-CoV-2 replication in infected cells. We demonstrate that 2NSP23 nanobody mRNA encapsulated into lipid nanoparticles (LNP), hereby referred to as LNP-mRNA-2NSP23, is internalized into HEK293 cells where it is translated. Using a combination of transcriptional profiling by RNA deep sequencing and qPCR analysis, we show that LNP-mRNA-2NSP23 protects HEK293-ACE2 cells and suppresses replication of several SARS-CoV-2 variants. Since, nanobody 2NSP23 specifically binds to viral Nsp9 (Esposito et al 2021), these observations strongly suggest that upon translation, the nanobody 2NSP23 inhibits viral replication by targeting Nsp9 in living cells. Given the conservation of Nsp9 across coronaviruses, we speculate that LNP-mRNA-2NSP23 may be translated into an innovative technology to generate a pan-coronavirus antiviral drug.

## Results

### The anti-Nsp9 2NSP23 nanobody mRNA is delivered as LNP and expressed in HEK293T cells

To overcome the challenge of nanobody uptake into cells, we used the lipid nanoparticle (LNP) technology (Maurer et al., 2001; Semple et al., 2010) for intracellular delivery of 2NSP23 nanobody as mRNA. LNPs containing nanobody mRNA, i.e. LNP-mRNA-2NSP23 (prepared by *in vitro* transcription) or tdTomato mRNA (LNP-mRNA-tdTomato) used as a control, were produced by nano-assembly microfluidic mixing technology (Hassett et al., 2019; Wei et al., 2020) (Figure 1a). HEK293T cells were treated with LNP-mRNA-tdTomato or LNP-mRNA-2NSP23 for 24 hrs and monitored with the Incucyte live-imaging technology or fluorescent microscope upon immunostaining, respectively (Figure 1b). Dilutions from 1/1000 to 1/25 of LNPs were used and red fluorescent signal of expressed tdTomato was analyzed 16h post treatment (Figure 1c). We observed an increasing number of HEK cells expressing tdTomato with a peak at 1/50 dilution. Imaging of the 1/50 LNP-mRNA-tdTomato-treated cells showed variability in the expression intensity of the protein (Figure 1d). For LNP-mRNA-2NSP23, we tested two different batches (S53 and S55) both at 1/50 dilution for 16hrs on HEK cells and stained the Nanobody proteins using a conjugated anti-alpaca antibody (Figure 1e). We observed batch to batch variability but both LNP-mRNA-2NSP23 were able to efficiently deliver the mRNA translated into protein in HEK293T cells. The analysis of the fluorescent signal intensity showed that batch S53 was statistically more efficient at delivering the Nanobody mRNA (Figure 1f). We conclude that the LNP technology is a suitable delivery method for nanobody mRNA cellular uptake and can be used as a general strategy for intracellular targeting of Nsp9 in cells infected with SARS-CoV-2.

**Figure 1.**
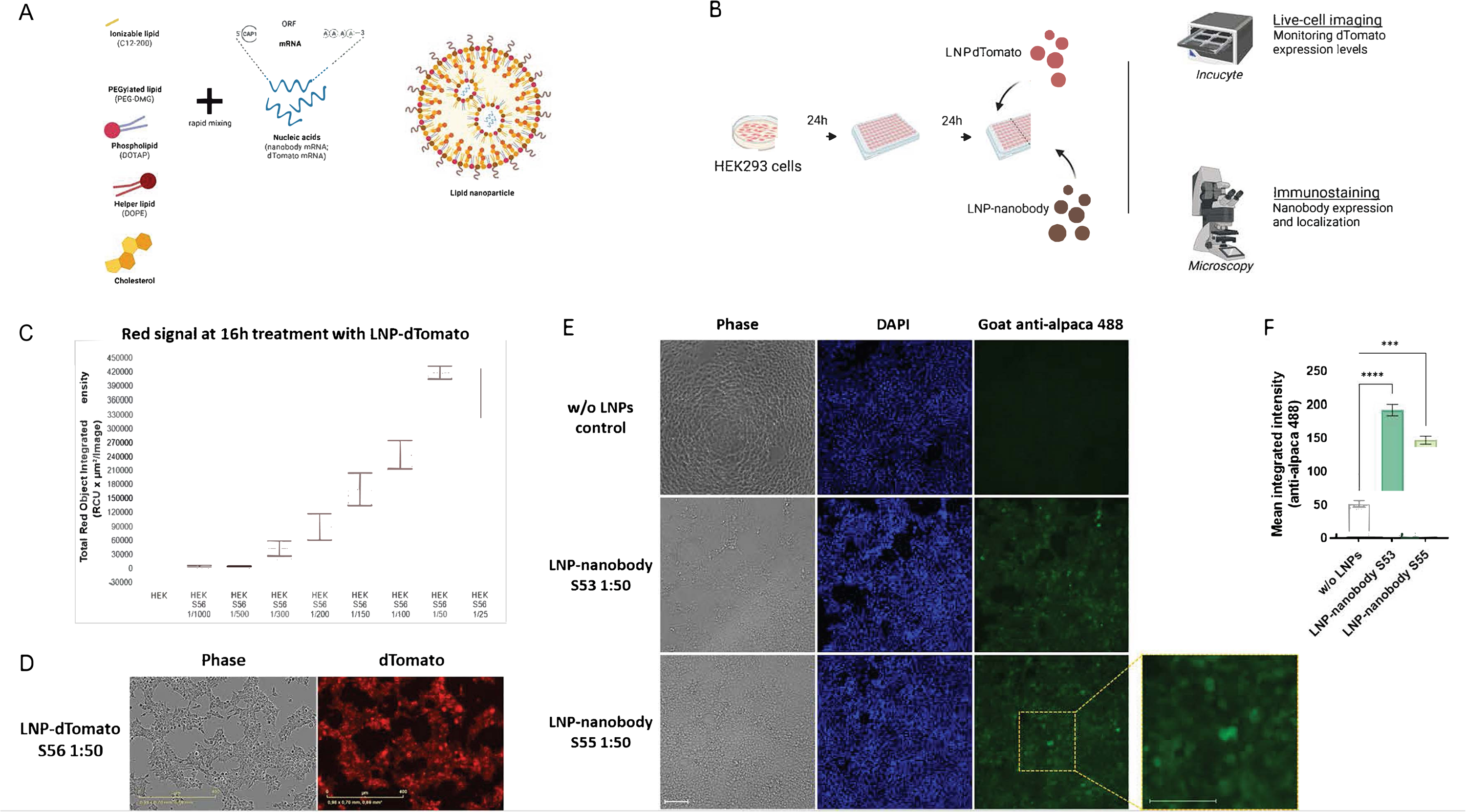
a) LNP production scheme. b) Protocol scheme of LNP treatment and readouts done for the HEK 293T cells. c) Intensity of red signal by the Incucyte at 16h treatment with different dilutions of LNP-dTomato. d) Representative pictures by the Incucyte at 16h treatment with LNP- dTomato S56 1:50. e) IF staining of the nanobody at 16h treatment with LNP-nanobody S53 1:50 and S55 1:50. f) Quantification of the IF staining.

### Intracellular LNP-mRNA-2NSP23 expression leads to SARS-CoV-2 inhibition

To evaluate the ability of SARS-CoV-2 Nsp9 protein-specific 2NSP23 nanobody to affect viral replication, we established a pipeline for testing LNP-mRNA-2NSP23 with different live infectious coronaviruses (Figure 2a). VeroE6 cells treated with serial dilutions of LNP-mRNA-2NSP23 or LNP-mRNA-tdTomato were first infected by an infectious clone (icSARS-CoV-2) that expresses the mNeonGreen as a reporter (Xie et al., 2020). The results showed that treatment with LNP-mRNA- 2NSP23 remarkably inhibited viral replication in a dose-dependent manner. Using the variable slope model in GraphPad Prism software, we calculated the concentration of LNP-mRNA-2NSP23 at which the green fluorescence intensity, reflecting viral replication, is reduced to half of its maximal value (EC50) and obtained 1.8 µM of LNP-mRNA-2NSP23 (Figure 2b), on par with previously obtained results using the FDA/EMA-approved antiviral small molecule Remdesivir (Kim et al., 2023). In contrast, LNP-mRNA-tdTomato treatment did not affect viral replication (Figure 2c and 2d), nor cell viability (cytotoxicity concentration (CC value) higher than 30 µM). We next used an independent recombinant infectious SARS-CoV-2 clone, harboring the nano-luciferase as a reporter marker (Hou et al., 2020). In two cell lines VeroE6 and HEK293-ACE2, we observed a dose-dependent reduction of relative nano-luciferase values when cells were treated with LNP-mRNA-2NSP23 targeting NSP9 protein (Figure 2e).

**Figure 2.**
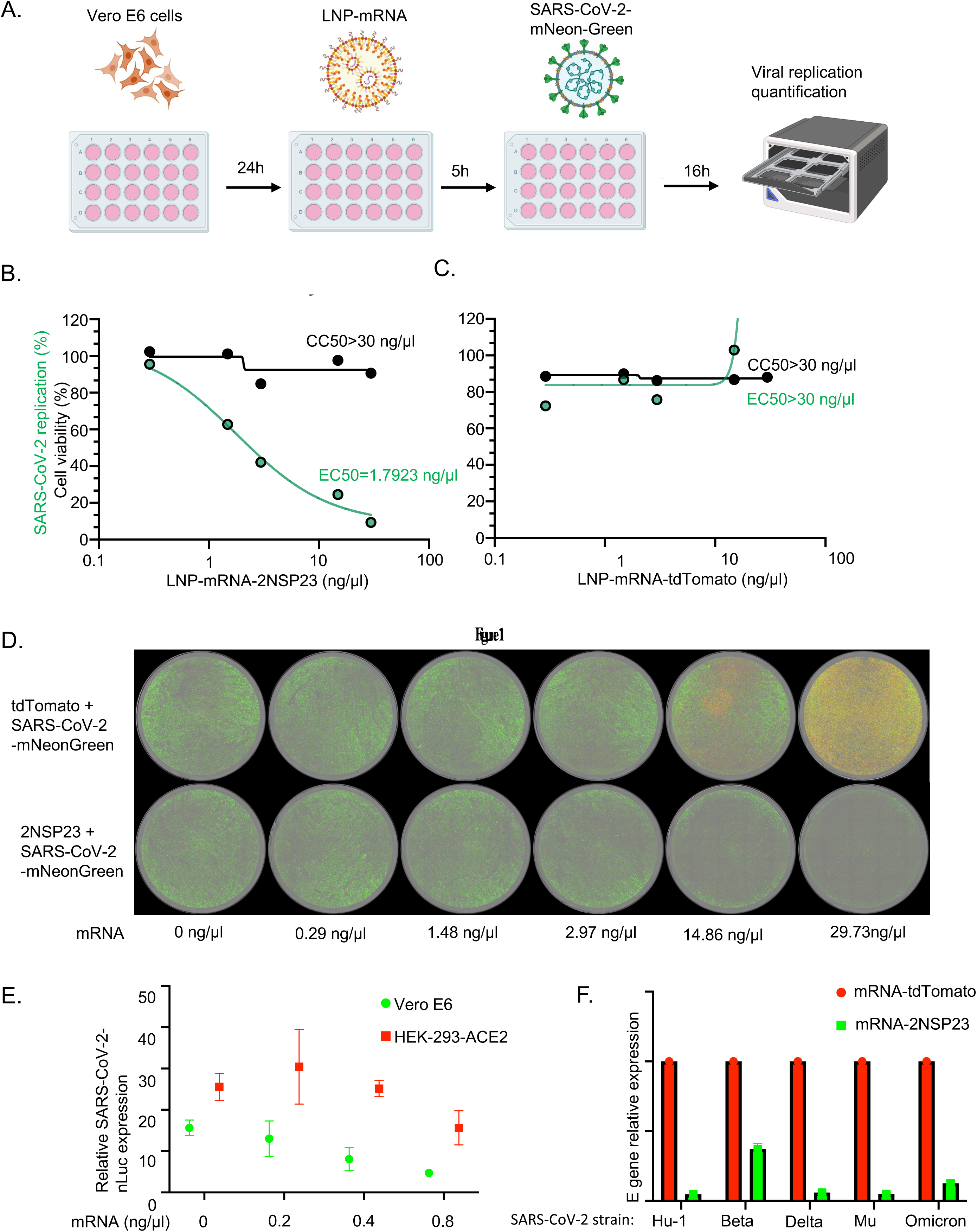
Inhibition of SARS-CoV-2 expression using mRNA encoding NSP9 nanobody. (A) Experimental pipeline. (B) Vero E6 (2× 10^5 cells/well), plated in a 24-well plate were treated with serial dilution with (29.7, 14.86, 2.97, 1.48, 0.29 ng/ul 0 of lipid nanoparticles (LNP)- encapsulated mRNA encoding a nanobody targeting NSP9. Cells were infected with SARS-CoV-2- mNeonGreen (0.1 of MOI), and Green fluorescence and cell viability were quantified using Incucyte S3 imaging. EC50 values were calculated via the variable slope model in GraphPad Prism v8. (C) As in B. but mRNA encoding a control dTomato fluorescent protein. (D) Representative images. The Green color corresponds to the expression of SARS-CoV-2-mNeonGreen. Yellow color indicates a concomitant expression of mNeonGreen and dTomato, respectively. (E) VeroE6 (green) or HEK293-ACE2 (red) plated in a 24-well plate were treated with indicated concentration of lipid nanoparticles (LNP)-encapsulated mRNA encoding a nanobody targeting NSP9, for 24 hours. Cells were infected with a SARS-CoV2-nLuc viral strain (0.01 of MOI), and luciferase quantified 24h post-infection. Nanoluciferase values are normalized using control cells treated with LNP-encapsulated mRNA encoding a tdTomato protein. (F) HEK293-ACE2 cells were seeded in 24-well plates for 24 hours, treated with 0.4 ng/ul of mRNA encoding a nanobody targeting NSP9 (green) or mRNA encoding dTomato (red) encapsulated in LNP, and infected with indicated viral strains at MOI of 0.01. Twenty-four hours post-infection, SARS-CoV-2 E gene was quantified by one-step qPCR from total RNA. Relative expression data, normalized to Ct values from cells treated with the control mRNA tdTomato are shown.

To find out if LNP-mRNA-2NSP23 treatment inhibits replication of different SARS-CoV-2 variants, HEK293-ACE2 cells treated with LNP-mRNA-2NSP23 or LNP-mRNA-tdTomato, respectively, were infected with the wild-type SARS-CoV-2 Wuhan strain and SARS-CoV-2 variants Beta, Delta, Mu and Omicron. RT-qPCR analysis of viral E gene expression shows that LNP-mRNA-2NSP23 (c= 0.4µg/ml) was able to abrogate viral replication at more than 90% for all variants except the SARS-CoV-2 variant Beta which was slightly less affected, with inhibition at ∼ 60% (Figure 2f) in comparison to LNP-mRNA-tdTomato treated samples.

Taken altogether, these results indicate that LNP-mRNA-2NSP23 is delivered into the cell, where it is translated into a fully functional nanobody that targets SARS-CoV-2 Nsp9 and inhibits viral replication.

### 2NSP23 nanobody treatment blocks viral replication of SARS-CoV-2 variants

While facilitating viral replication by contributing to RTC assembly, SARS-CoV-2 non-structural proteins, including Nsp9 and Nsp14, directly affect the host cell transcriptome (Zaffagni et al., 2022). In this way SARS-CoV-2 has a direct effect on the transcriptional profile of infected cells (Blanco-Melo et al., 2020; Chakraborty et al., 2021; Sun et al., 2021; Wyler et al., 2021). We, therefore, tested if treatment of HEK293T-ACE2 cells with the 2NSP23 nanobody rescues changes in host cell gene expression due to SARS-CoV-2 infection. Cells treated with LNP-mRNA-2NSP23 or LNP-mRNA-tdTomato were infected with SARS-CoV-2 variants UK (Beta), B1.617.X (Delta), Omicron and B1.621 (Mu) and total RNA was isolated from both infected and non-infected cells with or without LNP-mRNA-2NSP23 or LNP-mRNA-tdTomato treatment and subjected to deep RNA-sequencing. The sequencing reads were then aligned to human and SARS-CoV-2 genomes to measure the overall contribution of SARS-CoV-2 reads to total reads (Figure 3a). As expected, non-infected cells did not contain any SARS-CoV-2 RNA reads while in infected cells SARS-CoV-2 reads contributed to approximately 25 – 50% of total reads depending on the SARS-CoV-2 variant in control LNP-mRNA-tdTomato treated cells. In contrast, LNP-mRNA-2NSP23 treatment significantly reduced the SARS-CoV-2 reads to total reads ratio in all variants, strongly suggesting that SARS-CoV-2 RNA replication is inhibited upon nanobody treatment. Next, we performed hierarchical clustering based on similarity of all SARS-CoV-2 positive and negative samples treated with either LNP-mRNA-2NSP23 or LNP-mRNA-tdTomato (Figure 3b). We found that all LNP-mRNA-2NSP23 treated samples, regardless of their infection status, randomly clustered, while LNP-mRNA-tdTomato treated samples were separated based on SARS-CoV-2 positivity which indicated that LNP-mRNA-2NSP23 treatment suppresses the transcriptional effect of SARS-CoV-2 infection (Figure 3b). Next, we performed hierarchical clustering on RNA reads obtained from samples infected with SARS-CoV-2 UK variant treated with either LNP-mRNA-2NSP23 or LNP-mRNA-tdTomato as this variant showed the highest impact on transcriptional profiling and there is a clear separation between the SARS-CoV-2 infected dTomato treated cells in comparison to the rest of the cells (Figure 3c). Interestingly, this clustering is not seen in the principal component analysis, where samples are clearly separated between the treatments and infection status (Figure 3d). However, even here the variance between infected and non-infected cells in LNP-mRNA-2NSP23 treated cells is lower than LNP-mRNA-tdTomato-treated cells. The big variance between LNP-mRNA-2NSP23 and LNP-mRNA-tdTomato-treated cells suggests that expression of the 2NSP23 nanobody in cells causes a specific transcriptional response in cells (Figure 3d). To understand these differences, we performed pairwise comparisons between the 4 conditions to identify differentially expressed genes (Figure 3e-h). The MA plots obtained from this analysis show statistically significant differentially expressed (DE) genes (in red) which are either upregulated (upper part) or downregulated (lower part) in the first condition in comparison to the second condition. Figure 3e shows the effect of SARS-CoV-2 infection on control LNP-mRNA-tdTomato-treated cells with 2862 up-and 2668 down-regulated genes. In contrast, SARS-CoV-2 infection has only marginal effect on LNP-mRNA-2NSP23 treated cells with only 219 up-and 241 down-regulated genes (Figure 3f). The comparison between SARS-CoV-2 infected cells treated with LNP-mRNA-2NSP23 or LNP-mRNA-tdTomato shows similar pattern and number of DE genes (2003 up-and 2023 down-regulated) as seen between infected and non-infected LNP-mRNA-tdTomato treated samples suggesting that LNP-mRNA-2NSP23 expression in the cells suppresses the transcriptional outcome of viral infection to almost non-infected levels (Figure 3g). Finally, the comparison between the treatments in non-infected cells show the DE genes whose expression directly depends on LNP-mRNA-2NSP23 treatment with 460 genes to be upregulated and 271 genes to be downregulated (Figure 1h). In conclusion, transcriptional profiling of SARS-CoV-2 infected cells treated with either LNP-mRNA-2NSP23 or LNP-mRNA-tdTomato suggests specific role of anti-NSP9 antibodies in the reduction of viral replication.

**Figure 3.**
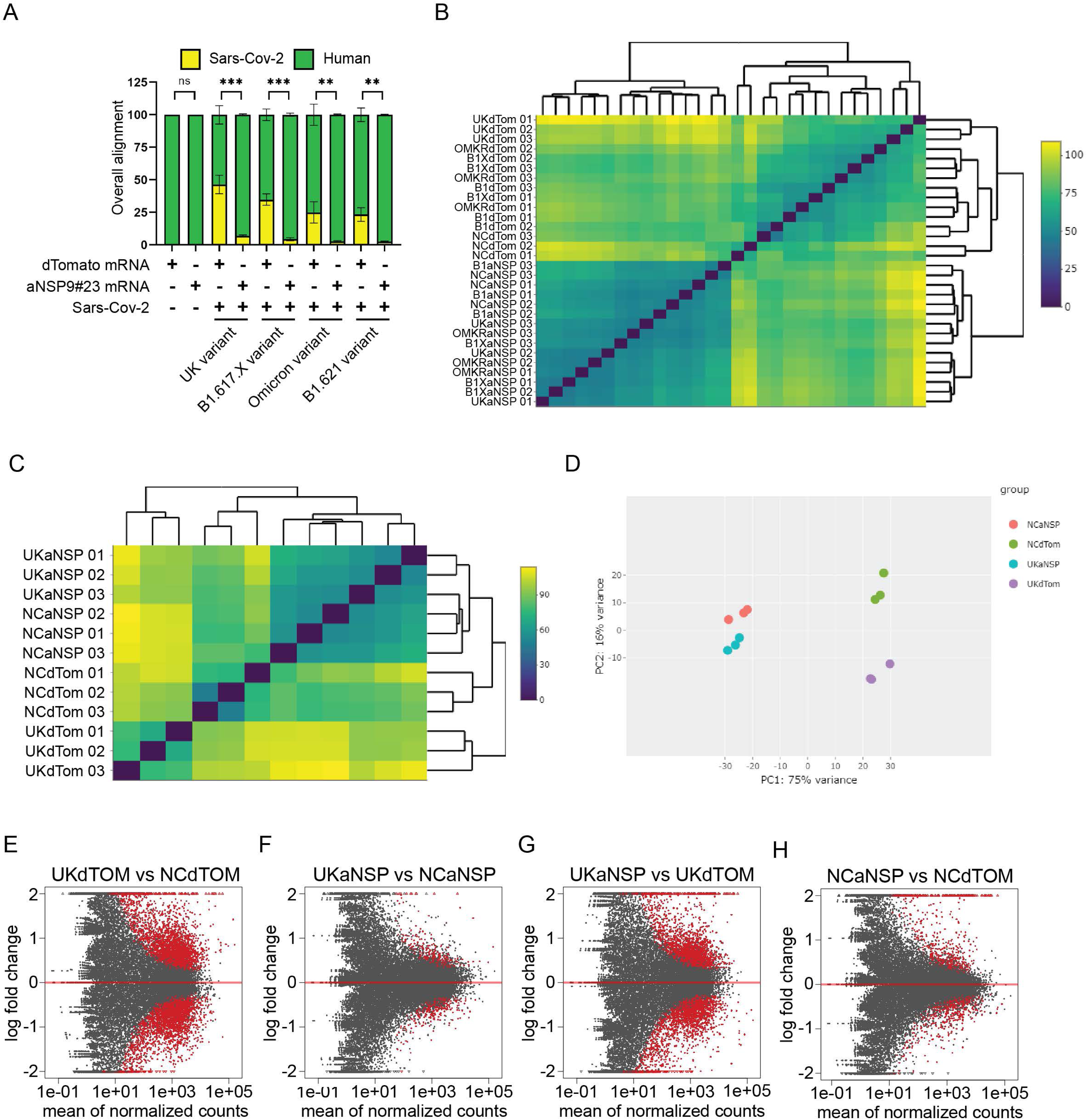
Nanobody treatment leads to a specific differential gene expression and suppression of SARS-CoV-2 replication (A) Overall alignment of sequencing reads to SARS-CoV-2 and human genomes in samples treated with anti-NSP9 nanobody mRNA or dTomato mRNA in non-infected or SARS-CoV-2 infected cells. (B) A heatmap representation of the distance matrix clustering based on similarity between all experimental samples. (C) A heatmap representation of the distance matrix clustering based on similarity between non-infected and SARS-CoV-2 UK variant-infected cells treated either with anti-NSP9 antibody mRNA or dTomato mRNA. (D) Principal Component Analysis (PCA) plot of non-infected and SARS-CoV-2 UK variant-infected cells treated either with anti-NSP9 antibody mRNA or dTomato mRNA (SARS-CoV-2 UK variant-infected cells treated with nanobodies - UKaNSP, cyan; or dTomato - UKdTOM, violet; and uninfected cells with nanobody expression - NCaNSP, red; or dTomato expression - NCdTOM, green). (E) MA plot showing differential gene expression between SARS-CoV-2 infected (UKdTOM) and non-infected (NCdTOM) cells without nanobody expression. Each red dot represents a single differentially expressed gene with upregulated genes in the upper part of the MA lot and downregulated genes in lower part of the MA plot. (F) MA plot showing differential gene expression between SARS-CoV-2 infected (UKaNSP) and non-infected (NCaNSP) cells with expression of anti-NSP9 nanobody. Each red dot represents a single differentially expressed gene with upregulated genes in the upper part of the MA lot and downregulated genes in lower part of the MA plot. (G) MA plot showing differential gene expression between SARS-CoV-2 infected cells with expression of anti-NSP9 nanobody (UKaNSP) or control dTomato (UKdTOM). Each red dot represents a single differentially expressed gene with upregulated genes in the upper part of the MA lot and downregulated genes in lower part of the MA plot. (H) MA plot showing differential gene expression between non-infected cells with expression of anti-NSP9 nanobody (NCaNSP) or control dTomato (NCdTOM). Each red dot represents a single differentially expressed gene with upregulated genes in the upper part of the MA lot and downregulated genes in lower part of the MA plot.

### 2NSP23 nanobody-treatment rescues mitochondrial function and activates antiviral immune response

During infection, the virus hijacks the cellular transcription machinery and mitochondria-suppressing innate immune response to produce viral proteins (Bhowal et al., 2023; Gatti et al., 2020; Shang et al., 2021; Singh et al., 2020). We performed gene ontology (GO) analysis of all differentially expressed genes between SARS-CoV-2 infected and non-infected cells treated with control LNP-mRNA-tdTomato and focused on the top hits in “biological process”, “cellular component” and “KEGG pathway” GO terms to identify the most affected cellular processes, compartments and pathways (Figure 4a). With almost no exceptions, the majority of GO terms in all three categories are associated with mitochondrial biogenesis and metabolism as well as with organization of chromatin structure and transcriptional regulation. To examine whether the affected pathways are activated or suppressed, we performed GO analysis on upregulated or downregulated genes separately (Figure 4b and 4c). Here, the GO terms associated with regulation of transcription are heavily upregulated (Figure 4b) while GO terms associated with mitochondrial function and oxidative phosphorylation are suppressed (Figure 4c), suggesting that SARS-CoV-2 virus fully hijacked cells for its own replication and spreading. As LNP-mRNA-2NSP23 treatment heavily reduced the number of differentially expressed genes between SARS-CoV-2 infected and non-infected cells, we next studied whether SARS-CoV-2 affects the same biological processes following intracellular nanobody expression. We first compared the overlap between the two data sets and found that the majority of differentially expressed genes observed in LNP-mRNA-2NSP23-treated samples is differentially expressed in LNP-mRNA-tdTomato-treated samples upon SARS-CoV-2 infection as well (Figure 4d). We performed the same GO analysis for these genes and found that both regulation of transcription and mitochondrial function are affected in LNP-mRNA-2NSP23-treated samples but to a significantly lower extent (Supplementary figure 1a-c). To see how nanobody treatment affects expression of individual genes, we took all differentially expressed genes between SARS-CoV-2 infected and non-infected cells treated with LNP-mRNA-tdTomato and plotted normalized read counts for each gene and each experimental condition in descending order in the form of heatmap (Figure 4e-g). The results show that SARS-CoV-2 infection has a prominent effect on expression profiles in LNP-mRNA-tdTomato-treated cells while treatment with LNP-mRNA-2NSP23 reduces not only the number of differentially expressed genes but also the level of expression of those genes which are significantly changed (Figure 4e). This can be seen more profoundly in selected groups of genes related to regulation of transcription (Figure 4f) and mitochondrial function (Figure 4g) where nanobody-treatment suppresses the negative effect of SARS-CoV-2 infection on cells. We, therefore, conclude that intracellular expression of 2NSP23 heavily rescues host cell gene expression programs that are otherwise dysregulated upon SARS-CoV-2 infection.

**Figure 4.**
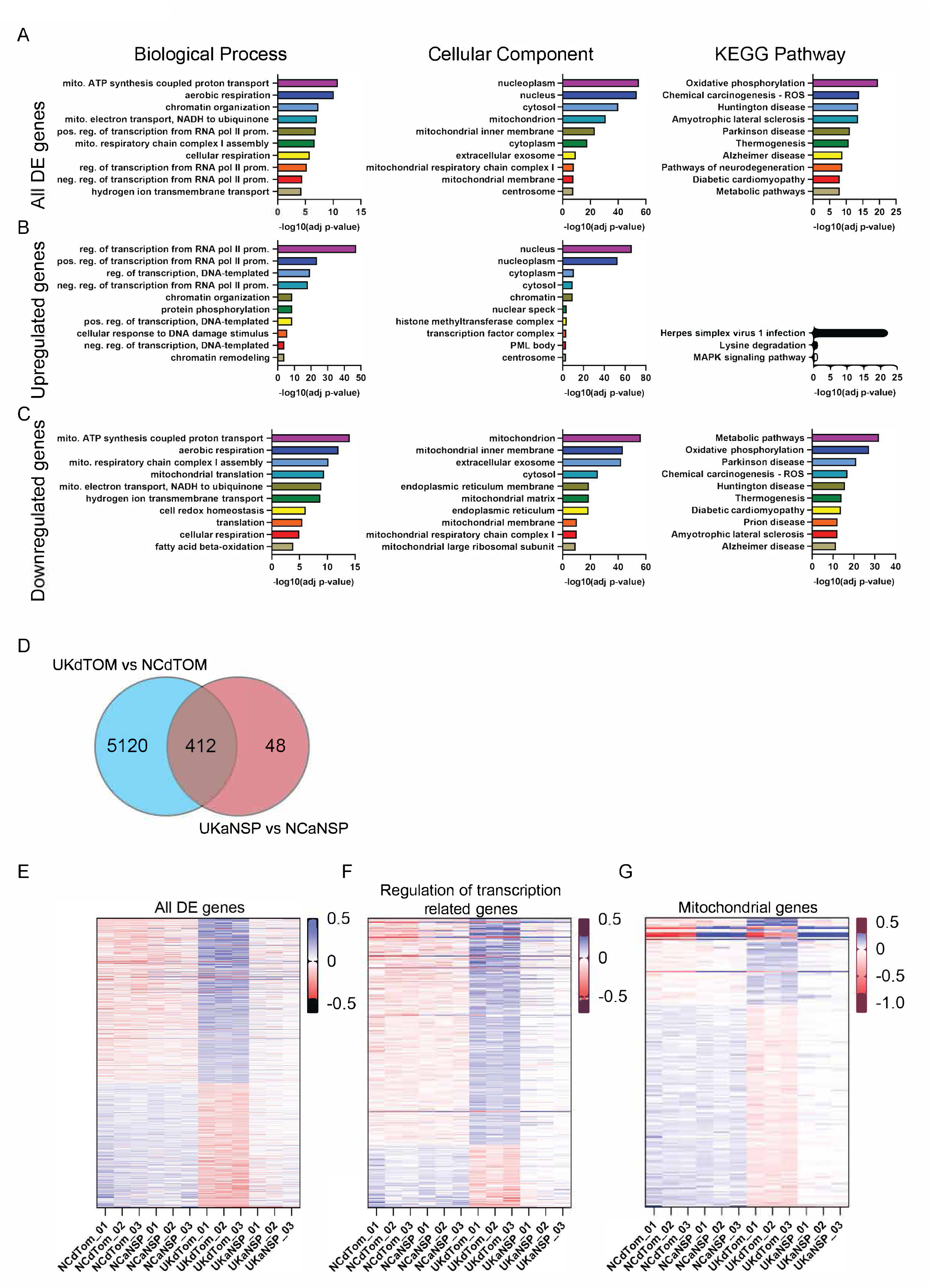
SARS-CoV-2 virus infection leads to differential gene expression of RNA Polymerase II transcription-regulatory genes and mitochondrial genes. (A) Top 10 GO terms associated with “biological process”, “cellular component” and “KEGG pathway” groups based on the analysis of all DE genes between infected and uninfected cells treated with dTomato (UKdTOM vs. NCdTOM). (B) Top GO terms associated with “biological process”, “cellular component” and “KEGG pathway” groups based on the analysis of upregulated genes in infected cells treated with dTomato in comparison to uninfected cells treated with dTomato. (C) Top GO terms associated with “biological process”, “cellular component” and “KEGG pathway” groups based on the analysis of downregulated genes in infected cells treated with dTomato in comparison to uninfected cells treated with dTomato (D) Venn diagram showing the number of specific and common genes which are differentially expressed upon SARS-CoV-2 infection in cells treated with dTomato (UKdTOM vs NCdTOM) and SARS-CoV-2 infection in cells treated with anti-NSP9 nanobody (UKaNSP vs NCaNSP). (E) Gene expression heatmap of normalized counts for each DE gene between infected and uninfected cells treated with dTomato (UKdTOM vs. NCdTOM) across each sample replicate. (F) Gene expression heatmap of normalized counts for each DE gene associated with GO term “Regulation of Pol II transcription” between infected and uninfected cells treated with dTomato (UKdTOM vs. NCdTOM) across each sample replicate. (G) Gene expression heatmap of normalized counts for each DE gene associated with GO term “Mitochondria” between infected and uninfected cells treated with dTomato (UKdTOM vs. NCdTOM) across each sample replicate.

We next explored which pathways are affected following intracellular 2NSP23 nanobody expression in both healthy and infected cells. For this purpose, we performed GO analysis of all differentially expressed genes between LNP-mRNA-2NSP23- and LNP-mRNA-tdTomato-treated non-infected cells and found multiple biological processes (BP) pertinent to antiviral immune response to be over-represented, including defense response to virus, negative regulation of viral genome replication, and innate immune response as well as several KEGG pathways associated with virus infection (Figure 5a). To see whether these processes are activated or suppressed in these cells, we performed GO analysis of upregulated and downregulated genes separately. As the majority of differentially expressed genes between LNP-mRNA-2NSP23- and LNP-mRNA-tdTomato-treated cells is upregulated, we did not get any significant GO terms for downregulated genes and GO terms associated with only upregulated genes copies the patterns seen in global DE analysis (Figure 5b). This indicates that introducing nanobodies into cells triggers the host immune response even prior to SARS-CoV-2 infection. To test whether immune system activation by nanobody treatment is persistent even after SARS-CoV-2 infection, we compared the differentially expressed genes between LNP-mRNA-2NSP23 and LNP-mRNA-tdTomato treatments in non-infected versus SARS-CoV-2 infected cells. We found that LNP-mRNA-2NSP23 treatment causes a very specific cellular response which is maintained and even strengthened upon virus infection (Figure 5c and 5d). Finally, to understand how the nanobody may activate an immune response, we compared the two most significant GO terms “defense response to virus” and “innate immune response” (Figure 5e) and plotted expression profiles of each single gene based on their specificity and expression pattern to common genes found in both groups (Figure 5f), genes found only in “defense response to virus” group (Figure 5g) and “innate immune response” specific group of genes (Figure 5h). While there is a subset of downregulated genes upon LNP-mRNA-2NSP23 treatment (Figure 5d), all differentially expressed genes related either to defense response to virus or innate immune response are upregulated suggesting very specific activation of antivirus response in 2NSP23 nanobody expressing cells. A remarkable observation is that treatment of cells with LNP-mRNA-2NSP23 elicited enrichment of genes related to defense response to viruses in uninfected cells. A number of genes in immune signaling pathways were heavily up-regulated in these healthy samples. For example, RSAD2, OAS1, OAS2, all of which being crucial components of interferon signaling pathways, were also found to be similarly affected by SARS-CoV-2 infection (Zhao et al., 2022), and their expression profiles in these treated samples resemble those in the infected cells (Figure 5f). Taken altogether these findings suggest that that 2NSP23 protects cells from SARS-CoV-2 infection by triggering genes involved in antiviral immune response.

**Figure 5.**
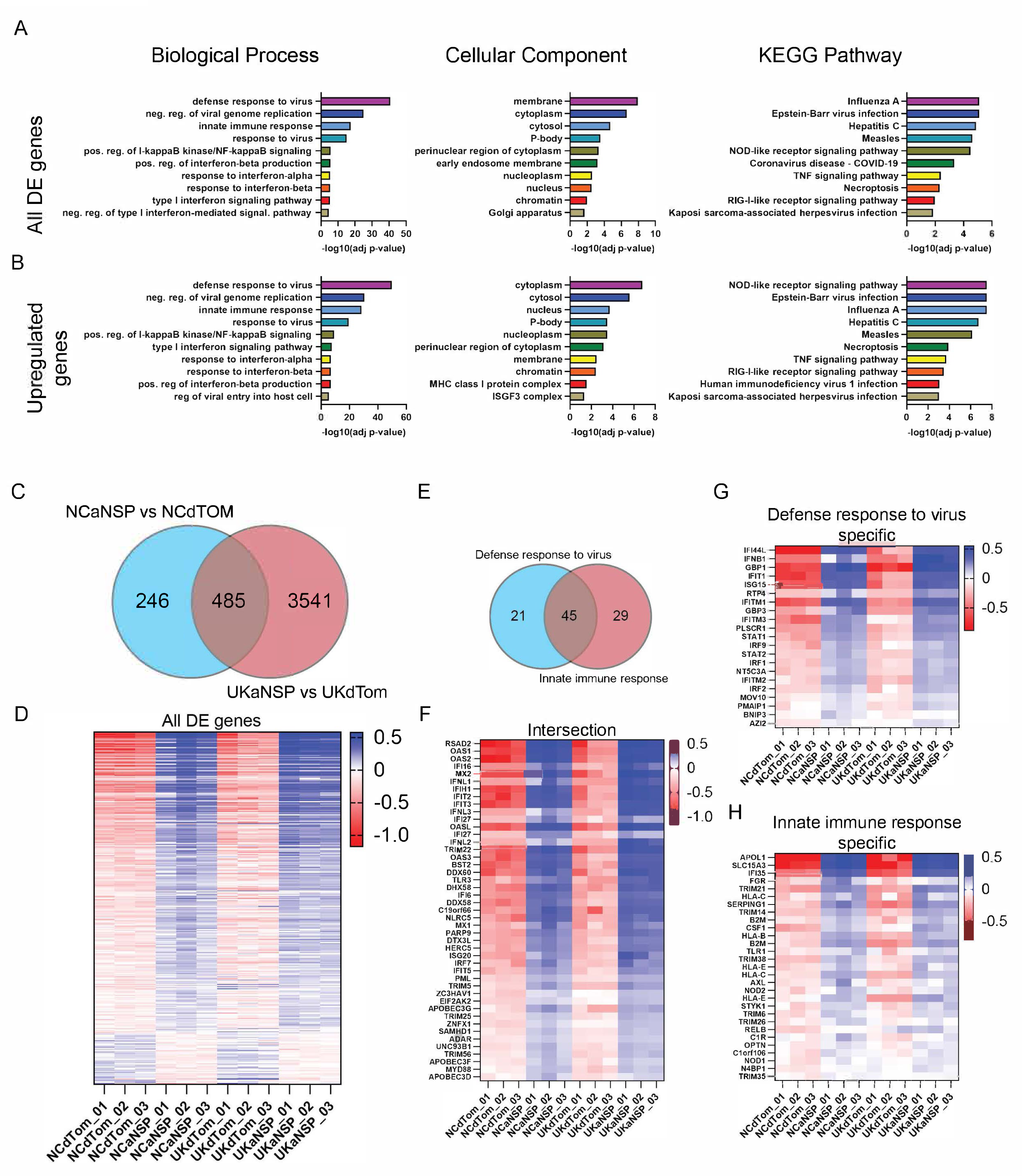
Nanobody treatment induces changes in gene expression related to host antiviral immune response regardless the SARS-CoV-2 infection. (A) Top 10 GO terms associated with “biological process”, “cellular component” and “KEGG pathway” groups based on the analysis of all DE genes between non-infected cells treated either with anti-NSP9 antibody mRNA or dTomato mRNA (NCaNSP vs. NCdTOM). (B) Top GO terms associated with “biological process”, “cellular component” and “KEGG pathway” groups based on the analysis of all upregulated genes in non-infected cells treated with antiNSP9 antibody in comparison to non-infected cells treated with dTomato. (C) Venn diagram showing the number of specific and common genes which are differentially expressed in nanobody treated non-infected cells and nanobody treated SARS-CoV- 2 infected cells in comparison to their respective dTomato-expressing control cells (NCaNSP vs NCdTOM and UKaNSP vs UKdTOM respectively). (D) Gene expression heatmap of normalized counts for each DE gene between anti-NSP9 antibody-treated and dTomato-treated non-infected cells (NCaNSP vs NCdTOM) across each sample replicate. (E) Venn diagram showing the intersection of differentially expressed genes associated with GO terms “Defense response to virus” and “Innate immune response” in non-infected cells treated with anti-NSP9 antibody or dTomato. (F) Gene expression heatmap of normalized counts for each DE gene found in intersection between GO terms “Defense response to virus” and “Innate immune response” in non-infected cells treated with anti-NSP9 antibody or dTomato. (G) Gene expression heatmap of normalized counts for each DE gene specific for GO term “Defense response to virus” in non-infected cells treated with anti-NSP9 antibody or dTomato. (H) Gene expression heatmap of normalized counts for each DE gene specific for GO term “Innate immune response” in non-infected cells treated with anti-NSP9 antibody or dTomato.

## Discussion

In this study, we have focused on the SARS-CoV-2 non-structural protein Nsp9 as a potential intracellular target to design compounds that might block replication of coronaviruses. Specifically, we have established a pipeline for intracellular expression of 2NSP23, a novel nanobody that specifically interacts with both recombinantly expressed Nsp9 and endogenous one in saliva of Covid-19 patients (Esposito et al., 2021). Earlier work by NMR spectroscopy showed that 2NSP23 stabilizes a short oligomeric form of Nsp9 (Esposito et al., 2021). As Nsp9 has been suggested to function in a monomeric/dimeric form to facilitate assembly of the RTC complex, we speculated that targeting viral replication using 2NSP23 nanobody, could serve as inhibitor of viral replication that would affect assembly of the RTC complex. To prove this hypothesis, we combined the use of 2NSP23 with RNA technology and lipid nanoparticles. Using this approach, we show that 2NSP23 is efficiently delivered into HEK293T cells in the form of mRNA encapsulated into LNPs. Following delivery, the 2NSP23 mRNA is intracellularly translated and blocks SARS-CoV-2 infection in cells. Analysis of RNA-Seq reads obtained from total RNA isolated from cells expressing the 2NSP23 nanobody and subjected to SARS-CoV-2 infection demonstrate the lack of viral RNA reads indicating that replication of SARS-CoV-2 RNA is inhibited when Nsp9 is targeted by the nanobody. Nsp9 is central for assembly of RTC complexes in SARS-CoV-2 and this is an essential step in viral replication. As endogenous Nsp9 seems to be quantitatively bound by 2NSP23, a likely mechanism of action leading to inhibition of viral RNA replication is a consequence of impaired RTC assembly due to the lack of Nsp9 availability.

Results from RNA-Seq experiments also indicate that in infected cells, 2NSP23 intracellular expression leads to a general rescue of the host cell transcriptional profile comparable to that of uninfected cells. This is particularly interesting considering the emerging view that viral proteins function in an orchestrated way to efficiently impact on the host cell transcriptome after viral infection. For instance, expression of exogenous SARS-CoV-2 Nsp14 in cells has been shown to provoke a dramatic remodeling of the transcriptome similar to that observed after SARS-CoV-2 infection. Nsp14 seems to contribute to remodeling of the transcriptome through a yet unclear mechanism that leads to alteration in the splicing of a set of genes and increased expression of circRNAs, all linked to innate immunity (Zaffagni et al., 2022). Our present work goes in the same direction since results from RNA-Seq experiments show significant differential gene expression upon SARS-CoV-2 infection, especially of those genes involved in transcriptional regulation and mitochondrial functions. Mitochondrion appeared to be the most impacted cellular component with metabolic pathways, especially oxidative phosphorylation, strongly down regulated. This is consistent with previous studies showing that SARS-CoV-2 viral damage causes and intensifies oxidative stress (Burtscher et al., 2020). More importantly, these mitochondrial genes involved in oxidative phosphorylation normally affected in infected cells were among those gene programs which were rescued in the presence of anti-Nsp9 nanobodies. Some of these genes include COX5B and NDUFS8 which are essential for the electron transport chain, and MRPL9, MRPL24, as well as other candidates, which are mitochondrial ribosomal proteins that govern not only oxidative phosphorylation, but also other cellular processes such as apoptosis and immune response. These results confirm that mitochondrial activity is a critical marker to examine if any treatment to the disease eventually works. It is remarkable that by expressing the anti-Nsp9 nanobody intracellularly, we are able to rescue these gene expression programs. In fact, these results suggest a potential role of Nsp9 as well as other Nsp proteins such as Nsp14 not only in RTC assembly but also in altering expression of cellular genes involved in essential host cell functions. How this is achieved remains to be further understood. However, given that 2NSP23 expression rescues these gene programs, it is likely that Nsp9, being an RNA-binding protein, might interact with nuclear factors and directly affect gene expression at transcriptional level in the host cells. So, 2NSP23 nanobody is ideally suited to function as potential antiviral to combat SARS-CoV-2 infection and infections by closely related viruses such as MERS and SARS-CoV considering the extremely low degree of mutagenicity of Nsp9 (Abbasian et al 2023). However, the remarkable finding that treatment of uninfected cells with 2NSP23 elicits enrichment of genes related to defense response to viruses also suggest a potentially much broader use of this nanobody in general prophylaxis to combat pathogens. Although the mechanisms underlying this specific event remain to be understood, we speculate that by triggering expression of genes involved in antiviral immune response, 2NSP23 protects cells from infection and provides broadly neutralizing activity against coronaviruses and other pathogens.

In summary, engineering modified mRNA molecules encoding highly potent nanobodies directed against key components of the SARS-CoV-2 RTC components such as Nsp9 is an attractive strategy to inhibit SARS-CoV-2 RNA replication. It is an efficient way of targeting intracellular viral proteins that are less prone to mutations in comparison to cell surface expressed proteins such as the Spike proteins and therefore, they may illuminate innovative solutions to combat pathogens. In addition, because of the larger interface area between the nanobody and its target, it is also anticipated that these mRNA-nanobodies-based therapies should drive fewer resistant mutants compared to small molecules targeting catalytic pockets for instance within Nsp12 enzyme. Translating those therapeutic mRNA into innovative antivirals should thus follow the same path as current mRNA vaccines against SARS-CoV-2.

## Methods

### Cell Culture

The HEK 293T cells (human embryonic kidney cells stably expressing the human ACE- 2 receptor) and Vero E6 cells (Epithelial cells an African green monkey, Cercopithecus aethiops) were grown in full Dulbecco’s Modified Eagle Medium (DMEM) containing 10% fetal bovine serum, 100 U/ml penicillin and 100 mg/ml streptomycin (Millipore-Sigma) in the humidified incubator at 37°C with 5% CO2.

### mRNA production

The 2NSP23 nanobody mRNA was prepared by in vitro translation of a DNA molecule of using the HiScribe™ T7 mRNA Kit with CleanCap® Reagent AG (New England BioLabs). The obtained mRNA has the following sequence and includes a 5’-CAP and a poly-A tail: aggaauugugagcggauaacaauucccucuagaauaauuuuguuuaacuuuaagaaggagauauaccaugggcaugcagCAGGUGCAGCUGCAGGAGUCUGGAGGAGGAUUGGUACAGCCUGGGGGCUCUCUGAGACUCUCCUGUGCAGCCUCUGGACUCGCCUUUAGUAUGUAUACCAUGGGCUGGUUCCGCCAGGCUCCAGGGAAGGAGCGUGAGUUUGUAGCAAUGAUUAUUUCAAGUGGUGAUAGCACCGACUACGCAGACUCCGUGAAGGGCCGAUUCACCAUCUCCAGGGACAACGGCAAGAACACGGUGUAUCUGCAAAUGGACAGCCUGAAACCUGAGGACACGGCCGUUUAUUACUGUGCAGCCCCAAAGUUUCGUUACUACUUUAGCACCUCUCCAGGUGAUUUUGAUUCCUGGGGCCAGGGGACCCAGGUCACCGUCUCCUCAGCGGCCGCAUACCCGUACGACGUUCCGGACUACGGUUCCCACCACCAUCACCAUCACUAGG

The nucleotide sequence of the corresponding DNA molecule encoding the 2NSP23 nanobody is: uaauacgacucacuauaggaggaauugugagcggauaacaauucccucuagaauaauuuuguuuaacuuuaagaaggagauauaccaugggcaugcagCAGGUGCAGCUGCAGGAGUCUGGAGGAGGAUUGGUACAGCCUGGGGGCUCUCUGAGACUCUCCUGUGCAGCCUCUGGACUCGCCUUUAGUAUGUAUACCAUGGGCUGGUUCCGCC AGGCUCCAGGGAAGGAGCGUGAGUUUGUAGCAAUGAUUAUUUCAAGUGGUGAUAGCACCGACUACGCAGACUCCGUGAAGGGCCGAUUCACCAUCUCCAGGGACAACGGCAAGAACACGGUGUAUCUGCAAAUGGACAGCCUGAAACCUGAGGACACGGCCGUUUAUUACUGUGCAGCCCCAAAGUUUCGUUACUACUUUAGCACCUCUCCAGGUGAUUUUGAUUCCUGGGGCCAGGGGACCCAGGUCACCGUCUCCUCAGCGGCCGCAUACCCGUACGACGUUCCGGACUACGGUUCCCACCACCAUCACCAUCACUAGGAAUUC

### LNP production

The lipid nanoparticle (LNP-mRNA) was prepared using NanoAssemblr® (Precision Nanosystems) microfluidic mixing technology under time invariant conditions. 2 ml of an aqueous solution containing the mRNA at a concentration of 174 µg/ml in aqueous 70 mM acetate buffer, pH 4.0, was mixed with 1 ml aqueous ethanolic lipid solution containing 12.5 mM lipids to form the nanoparticles. The flow rate ratio between the aqueous solution and the aqueous ethanolic lipid solution was 3:1, and the total flow rate was 12 ml/min.

The aqueous ethanolic lipid solution was prepared by dissolving C12-200 (Corden Pharma), DOPE (1,2-dioleoyl-sn-glycero-3-phosphoethanolamine), Cholesterol, DMG-PEG (PEGylated myristoyl diglyceride; Avanti catalog no. 880151P-1G) and DOTAP (1,2-di-(9Z-octadecenoyl)-3- trimethylammonium propane methylsulfate Avanti catalog no. 890890-200mg) in a molar ratio of 29.8 : 13.6 : 39.5 : 2.1 : 15 in ethanol. This was done by preparing separate 12.5 mM solutions of each lipid in the ethanol and mixing the solutions in the ratio given above to give the aqueous ethanolic lipid solution.

1.6 ml of the obtained LNP-mRNA product was immediately diluted with 64 ml PBS (1X) and concentrated to 1.5 ml at 2000g for 30 minutes at 20°C using Amicon® Ultra-15 centrifugal filtration tubes. Finally, the LNP sample was sterilized with 0.2 µm syringe filter and stored at 4°C. Size distribution of particles 70.7nm, particle number 5.31E+11 and zeta potential 0.51 mV were measured with Zetasizer Ultra (Malvern Panalytical Ltd).

### mRNA delivery with LNPs

HEK 293T cells were plated in complete medium at a density of 30’000 cells and 100 µl per well in a 96-well plate (transparent flat bottom, Sarstedt) and incubated for 24h in 5% CO2 at 37°C. LNPs carrying mRNA for dTomato or for the nanobody were added to the cells at different dilutions and the cells were incubated overnight (16h) at the 5% CO2 37°C incucyte incubator. Untreated HEK 293T cells were used as an auto-fluorescence control.

### Analysis of dTomato expression levels after LNP delivery by live-cell imaging

For monitoring dTomato expression levels, cells were scanned for red signal detection during 16h incubation with the different dilutions of LNP-dTomato. Live-cell imaging was done with a S3 Incucyte (S3/SX1 G/R Optical Module, Sartorius) for red and phase channels, acquisition time 400ms for both channels and magnification 10x. Total Red Object Integrated Intensity was quantified for each condition using the Incucyte Software Basic Analyser. The analysis definitions used for red signal were segmentation Top-Hat, radius 40µm and threshold 0.4 RCU with Edge Split Off. A filter for a minimum of 20 µm2 of area was used to remove debris from the analysis.

### Analysis of nanobody expression levels after LNP delivery by immunostaining

After 16h of incubation with the LNP-nanobody, cells were fixed by adding 100 µl 8% PFA for 20 min at RT (final concentration of PFA is 4%; PFA from VWR). All media was then gently removed and cells were washed 3 times for 5 minutes with 150 µl PBS. After the last wash, 100 µl of a blocking/permeabilization buffer (5% BSA and 0.2% Triton in PBS; BSA from Sigma; TritonX-100 from VWR) was added and cells were incubated for 1h at RT. Cells were then washed 3 times for 5 minutes with 150 µl PBS and staining was performed. For immunostaining, 50 µl of a 1:100 dilution of the AF488 goat anti-alpaca secondary antibody was used (Jackson Immunoresearch 0.25MG Alexa Fluor 488-AffiniPure Goat Anti-Alpaca IgG, VHH domain, from Fisher Scientific). For all cases, antibody was prepared fresh at 1:100 in PBA (1% BSA in PBS). Incubation with the antibody was performed for 1h at RT and protected from light. After nanobody staining cells were washed 3 times for 5 minutes with 150 µl PBS and then incubated for 5 minutes with 100 µl of DAPI 1:1000 in PBS for nuclear staining (DAPI from Sigma). The last 3 washes were then performed and cells were imaged directly at the 96-well plate with an inverted DMi8 microscope (Leica). For image acquisition the same exposure times were used for each correspondent channel, between the different conditions. Quantification of the AF488 goat anti-alpaca signal was performed with CellProfiler.

### SARS-CoV2 Infection assays

2×10^5 Vero E6 cells or HEK293-ACE-2 were cultured in 24-well plates in Dulbecco’s Modified Eagle medium (DMEM, Sigma D5796) containing 10% of fetal bovine serum (FBS), 2 mM of L-glutamine, 100 units of penicillin and 100 g/l of streptomycin in a cell culture incubator at 37 degrees with 5% CO2 and 95% humidity. After 24 h, cells were treated with increased concentration of mRNA molecules encoding for the nanobody NSP23 targeting NSP9 protein or mRNA molecules encoding dTomato, as a control. The mRNAs were encapsulated in lipid nanoparticles (LNP), with initial working concentrations of mRNA and LNP of 200 µg/ml and 5mM respectively, and 5 serial dilutions of two-fold were used. After 8 hours of incubation, cells were infected with indicated strains of SARS-CoV-2 at 0.1 of MOI. The next day, after 16 hours of culture, cells were either lysed for RNA extraction, nano-luciferase quantified in the case of icSARS-CoV2-nuLuc, or fixed with 3,7% paraformaldehyde during 20 min followed by fluorescent analysis using Incucyte S3 instrument. For the icSARS-CoV2-mNeonGreen replication efficiency, fluorescence quantification was performed using Incucyte software and EC50 values were calculated via the variable slope model in GraphPad Prism v8. For the icSARS-CoV2-nuLuc, luminescence was quantified using Molecular Devices Instrument. Cells viability was measured using the Cell Titer-Glo® Luminescent Cell Viability Assay kit (Promega, G7750). Nano-luminescence values were normalized to corresponding viability data.

### RNA extraction and RT-qPCR

HEK-293-ACE2 cells were infected as above with different strains of SARS-CoV-2 (Wuhan = alpha, UK = beta, B1.621 = mu, B1.617.x = delta, and Omicron=B1.1.529), and treated with LNP(10µM)-mRNA(0.4µg/ml) complexes encoding either the nanobody NSP23 targeting NSP9 or mRNA molecules encoding dTomato, as a control. Twenty-four hours post-infection, cells were washed with PBS and lysed with RNA extraction buffer (Nucleo spin RNA extraction kit, Macherey-Nagel, Cat# 740955). Total RNA was extracted according to the manufacturer instructions, concentrations determined using Nanodrop (Thermo), and the integrity of RNA verified using Agilent 2100 Bioanalyser Systems. One nanogram of RNA was subsequently used in a one-step RT-qPCR for the E gene of SARS-CoV-2 using the following primers: E_Sarbeco_F1: 5’- ACAGGTACGTTAATAGTTAATAGCGT-3’ and E_Sarbeco_R2: 5’-ATATTGCAGCAGTACGCACACA-3’ for amplification and the double dye probe E_Sarbeco_P1 5’-(FAM)ACACTAGCCATCCTTACTGCGCTTCG (BHQ)-3’ as previously described (Coupeau et al. Methods Protoc. 2020 Aug 18;3(3):59.doi: 10.3390/mps3030059.)

### RNA Sequencing Analysis

HEK 293-ACE2 cells either infected with the SARS-CoV-2 virus or uninfected, treated with LNPs containing mRNA sequences of either the anti-NSP9 nanobody or tdTomato were used for total mRNA isolation by TRI Reagent (Millipore-Sigma) according to the manufacturer’s protocol. Three biological replicates for each condition were used for transcriptional profiling. The RNA-Seq library was prepared by using the NEBNext Ultra II RNA Library Prep Kit for Illumina (NEB) and sequenced with the NextSeq 500/550 sequencing platform (performed at the NYUAD Sequencing Center). Raw FASTQ sequenced reads where first assessed for quality using FastQC v0.11.5 (https://www.bioinformatics.babraham.ac.uk/projects/fastqc/). The reads where then passed through Trimmomatic v0.36 for quality trimming and adapter sequence removal, with the parameters (ILLUMINACLIP: trimmomatic_adapter.fa:2:30:10 TRAILING:3 LEADING:3 SLIDINGWINDOW:4:15 MINLEN:36) (Bolger et al., 2014). The surviving trimmed read pairs were then processed with Fastp in order to remove poly-G tails and Novaseq/Nextseq specific artefacts (Chen et al., 2018). Following the quality trimming, the reads were assessed again using FastQC.

Post QC and QT, the reads were aligned to the human reference genome GRCh38.81 using HISAT2 (Kim et al., 2015) with the default parameters and additionally by providing the –dta flag. The resulting SAM alignments were then converted to BAM, whilst retaining only the unmapped read pairs using SAMtools (Li et al., 2009) v1.9 “samtools view -f 12 -F 256 -b INPUT.sam > OUTPUT.bam”. The resulting unmapped BAM reads were then sorted and converted to fastq reads using samtools sort ”samtools sort -@ 14 -o OUTPUT.sorted.bam INPUT.bam” and samtools fastq “samtools fastq -1 read1.fastq -2 read2.fastq -@ 14 INPUT.sorted.bam”. These unmapped reads where then aligned to the SARS-COV-2 reference genome available through NCBI with the accession NC_045512.2 using HISAT2 with the same parameters that were used in the initial mapping to the human reference. The resulting SAM alignments were then converted to BAM format and coordinate sorted using samtools view and samtools sort respectively. Read groups were then added to the sorted SARS-COV-2 mapped BAM files using Picard AddOrReplaceReadGroups (http://broadinstitute.github.io/picard/). A consensus genome was then generated for each SARS-COV-2 sample using samtools mpileup and ivar (Grubaugh et al., 2019) version 1.3 with the minimum depth set at 20 “samtools mpileup -A -d 0 -Q 0 INPUT.sorted.bam | ivar consensus -m 20 -p OUTPUT.sars-cov-2-consensus.fasta”. Finally, Qualimap (Garcia-Alcalde et al., 2012) v2.2.2 was used to generate alignment specific QC metrics per sample, both for alignments vs the human reference, as well as the SARS-COV-2 reference. The BAM alignment files were processed using HTseq-count, using the reference annotation file to produce raw counts for each sample. The raw counts were then analyzed using the online analysis portal NASQAR (http://nasqar.abudhabi.nyu.edu/), to merge, normalize, and identify differentially expressed genes (DEG). DEG (log2(FC) ≥ 0.5 and adjusted p-value of <0.05 for upregulated genes, and log2(FC) ≤ −0.5 and adjusted p-value of <0.05 for downregulated genes) between each sample combination were subjected to GO enrichment using DAVID Bioinformatics (https://david.ncifcrf.gov/) (Huang da et al., 2009). The full RNA-Seq data set was deposited inthe Gene Expression Omnibus (GEO) database under accession number GSE244714.

## Data Availability

Gene expression data are publicly available in the Gene Expression Omnibus (GEO) database under accession number GSE244714.

## Acknowledgments

This work is supported by grants from NYU Abu Dhabi, the Sheikh Hamdan Bin Rashid Al Maktoum Award for Medical Sciences, a donation from the Cipriani family, a Covid-19 Facilitator Grant from NYU Abu Dhabi and Tamkeen under the NYU Abu Dhabi Research Institute Award to the NYUAD Center for Genomics and Systems Biology (ADHPG-CGSB) to PP. We thank the NYU Abu Dhabi Center for Genomics and Systems Biology, in particular Marc Arnoux and Mehar Sultana for RNA sequencing. We appreciate the computational platform provided by the Center for Genomics and Systems Biology and the NYU Abu Dhabi HPC team and are especially thankful to Nizar Drou for technical help.

## Author Contributions Statement

PP conceived the research and wrote the manuscript with JCT. TV and SS performed all RNA-seq analyses, and data analysis and prepared the corresponding figures. JB and SBM performed functional assays and LE and CB performed the imaging. HPH and SD performed LNP characterization. PP, JCT, ML and GE analysed the results. PP supervised all the research. All authors read and approved the manuscript.

## Competing interests

PP, GE, HTH and SCD are part of a US patent filed by New York University in Abu Dhabi jointly with ISAR Biosciences. The other authors declare no competing interests.

---

**Supplementary figure 1:** The GO analysis of non-infected and SARS-CoV-2 infected cells treated with anti-NSP9 antibody leads to similar GO association as in dTomato-treated samples. (A) Top 10 GO terms associated with “biological process”, “cellular component” and “KEGG pathway” groups based on the analysis of all DE genes between infected and uninfected cells treated with anti-NSP9 antibody (UKdaNSP vs. NCaNSP). (B) Top GO terms associated with “biological process”, “cellular component” and “KEGG pathway” groups based on the analysis of all upregulated genes in infected cells treated with anti-NSP9 antibody in comparison to uninfected control cells. (C) Top GO terms associated with “biological process”, “cellular component” and “KEGG pathway” groups based on the analysis of all downregulated genes in infected cells treated with anti-NSP9 antibody in comparison to uninfected control cells.

## Notes

### Competing Interest Statement

The authors have declared no competing interest.

