## Supplementary figures and images for "Nanobodies against SARS-CoV-2 non-structural protein Nsp9 inhibit viral replication by targeting innate immunity"

### Supplementary figure 1

Supplementary figure 1

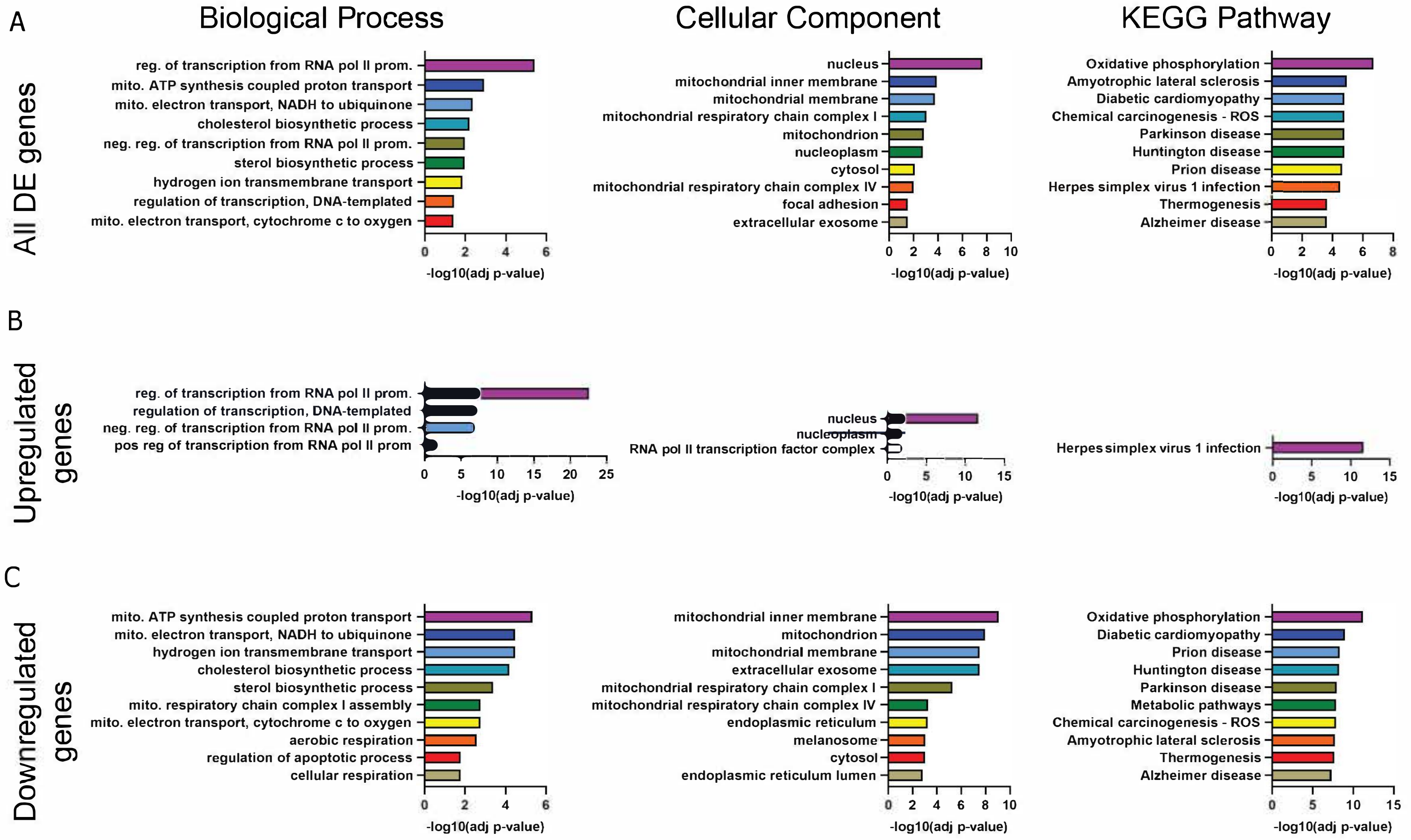
